# Zebrafish use conserved CLR and TLR signaling pathways to respond to fungal PAMPs in zymosan

**DOI:** 10.1101/2024.06.24.600417

**Authors:** Erin Glass, Stephan L. Robinson, Emily E. Rosowski

## Abstract

Pattern recognition receptors (PRRs) such as C-type lectin receptors (CLRs) and Toll-like receptors (TLRs) are used by hosts to recognize pathogen-associated molecular patterns (PAMPs) in microorganisms and to initiate innate immune responses. While PRRs exist across invertebrate and vertebrate species, the functional homology of many of these receptors is still unclear. In this study, we investigate the innate immune response of zebrafish larvae to zymosan, a β-glucan-containing particle derived from fungal cell walls. Macrophages and neutrophils robustly respond to zymosan and are required for zymosan-induced activation of the NF-κB transcription factor. Full activation of NF-κB in response to zymosan depends on Card9/Syk and Myd88, conserved CLR and TLR adaptor proteins, respectively. Two putative CLRs, Clec4c and Sclra, are both required for maximal sensing of zymosan and NF-κB activation. Altogether, we identify conserved PRRs and PRR signaling pathways in larval zebrafish that promote recognition of fungal PAMPs. These results inform modeling of human fungal infections in zebrafish and increase our knowledge of the evolution and conservation of PRR pathways in vertebrates.

## 1. Introduction

Zebrafish, and particularly larval zebrafish, have emerged as an exciting vertebrate host model for studying host-microbe interactions due to their low-cost, high fecundity, simple genetic techniques, and the ability to perform repeated, live, non-invasive imaging (Gomes and Mostowy, 2020, Stream and Madigan, 2022). In particular, this host model has been a source of new findings related to the pathogenesis of infections with human fungal pathogens, including *Aspergillus fumigatus*, *Candida albicans*, and *Cryptococcus neoformans* (Rosowski, Knox et al., 2018). To fully take advantage of zebrafish in modeling human fungal pathogen infection, we need to understand more about the similarities and differences between human and zebrafish immune mechanisms. In particular, a major gap in knowledge is the function of different innate immune receptors and pattern recognition receptors (PRRs) in zebrafish and other teleost fish (Li, Y. et al., 2017, Wcisel and Yoder, 2016, Zelensky and Gready, 2004).

Fungi possess a wide variety of pathogen-associated molecular patterns (PAMPs) in the cell wall, including β-glucan, mannans and mannosylated proteins, chitin, and melanin (Gow et al., 2017). These PAMPs are recognized by a large set of PRRs in mammals, including C-type lectin receptors (CLRs) and Toll-like receptors (TLRs) (Hatinguais et al., 2020). β-glucan is primarily detected by the CLR Dectin-1 (*CLEC7A* gene in mice) (Brown, G. D. and Gordon, 2001, Yokota et al., 2001), although other CLRs and non-CLR receptors can also recognize this sugar (Briard et al., 2019, de Jong, Marein A W P et al., 2010, Evans et al., 2005, Guo et al., 2018, Ross et al., 1987, Swidergall et al., 2018, Zani et al., 2015). Dectin-1-deficient mice are more susceptible to *A. fumigatus* (Werner et al., 2009) and some strains of *C. albicans* (Marakalala et al., 2013, Taylor, Philip R. et al., 2007), and Dectin-1 polymorphisms in humans are associated with increased risk of mucocutaneous fungal infections (Ferwerda et al., 2009) and invasive aspergillosis (Cunha et al., 2010, Sainz et al., 2012), underlining the role of this receptor in immunity against fungi. After recognition of β-glucan, Dectin-1 signals through the adaptor proteins Syk and Card9 to activate NF-κB transcription factors and expression of pro-inflammatory genes to control fungal infections (Hatinguais et al., 2023).

As in humans, immunocompetent larval zebrafish are largely resistant to infection with *A. fumigatus* and *C. albicans* (Gratacap et al., 2017, Knox et al., 2014), even before these fish have a fully competent adaptive immune system which develops after more than two weeks of age (Masud et al., 2017, Page et al., 2013). Innate immune cells, including neutrophils and macrophages, respond to these fungi to control the infection, demonstrating that zebrafish cells must have receptors that can sense PAMPs on these fungi. Several lines of evidence further demonstrate that zebrafish can respond to β-glucan specifically. First, exposure of zebrafish to β-glucan triggers both trained immunity (Darroch et al., 2022) and increased resistance to viral infection (Liang et al., 2022). Second, phagocytes in larval zebrafish can shuttle fungal spores from one cell to another and a β-glucan signal was sufficient to instigate this shuttling (Pazhakh et al., 2019). However, whether the zebrafish genome codes for a Dectin-1 homolog that recognizes β-glucan has remained unclear. After a failure to identify such a gene through sequence homology, Petit et al recently identified a genomic region in zebrafish that has partial conservation of synteny with the genomic location of Dectin-1 in humans and mouse that contains two putative CLRs (Petit et al., 2019).

In this study, we sought to further define the innate immune response to fungal PAMPs and to test the role of these two putative CLRs in this response. We demonstrate that zebrafish respond robustly to zymosan, a fungal-derived particle primarily made up of β-glucans but that also contains mannans (Di Carlo and Fiore, 1958).

Both macrophages and neutrophils are recruited to the site of zymosan injection, phagocytose zymosan particles, and activate the transcription factor NF-κB. The presence of phagocytes is required for NF-κB activation in response to zymosan, with macrophages playing a larger role than neutrophils. We find that zebrafish use both CLR and TLR signaling pathways to initiate NF-κB activation and identify roles for both previously identified putative CLRs (Sclra and Clec4c) and Tlr2. Overall, we further define the innate immune response to fungal PAMPs in larval zebrafish and find that two putative CLRs in the zebrafish genome have a role in responding to these ligands.

## 2. Materials and Methods

### 2.1 Zebrafish lines and husbandry

Adult and larval zebrafish were maintained and handled according to protocols approved by the Clemson University Institutional Animal Care and Use Committee (AUP2021-0109, AUP2022-0093, and AUP2022-0111) and following the Guide for the Care and Use of Laboratory Animals. Adult zebrafish were maintained at 28°C in a 14-/10-hour light/dark cycle. All mutant and transgenic fish lines used in this study are listed in Table 1 and were maintained in the AB background. Adults were spawned naturally, and embryos were kept in E3 medium with methylene blue at 28°C. Sex is not yet determined in zebrafish embryos and larvae. For all imaging experiments, embryos were treated with 200 µM N-phenylthiourea (PTU) from 24 hours post fertilization onwards. Embryos were manually dechorionated and anesthetized in 0.3 mg/mL buffered tricaine prior to any experimental manipulations. All mutant lines were maintained as heterozygotes by out-crossing to wild-type or transgenic lines.

**Table 1.** Zebrafish lines used in this study.

| Table 1. Zebrafish lines used in this study |  |  |
| --- | --- | --- |
| Line name | Purpose | Reference |
| Tg(*NF-κB RE::EGFP*) | Report NF-κB activation | (Kanter et al., 2011) |
| Tg(*lyz::BFP*) | Label neutrophil cytoplasm | (Rosowski, Raffa et al., 2018) |
| Tg(*mpeg1::mCherry-H2B*) | Label macrophage nuclei | (Vincent et al., 2016) |
| Tg(*mfap4::BFP*) | Label macrophage cytoplasm | (Thrikawala, Savini U. et al., 2023) |
| Tg(*mpx::mCherry-2A-rac2D57N*) | Neutrophil-defective | (Deng et al., 2011) |
| *irf8* st95 | Macrophage-deficient | (Shiau et al., 2015) |
| *card9* sa11441 | Point mutant causing a premature stop mutation in *card9* | Zebrafish Mutation Project, ZIRC |
| *clec4c* d7 | 7 bp deletion in *clec4c* predicted transmembrane domain | This study |

For genotyping, genomic DNA (gDNA) from individual embryos/larvae or from tail fin tissue was isolated in 50 mM NaOH at 95°C as previously described (Meeker et al., 2007). The *irf8* st95 mutation abolishes an AvaI restriction site and this line was genotyped by PCR (primers in Table 2) and AvaI digest (NEB), as previously described (Shiau et al., 2015). The *card9* sa11441 line was received from the Zebrafish International Resource Center (ZIRC) and this point mutation introduces a premature stop codon and abolishes an Hpy118I site and the line was genotyped by PCR (primers in Table 2) and Hpy118I digest (NEB) (Supp Fig 1B). All transgenic lines were screened for fluorescence by microscopy, as described below.

**Table 2.**
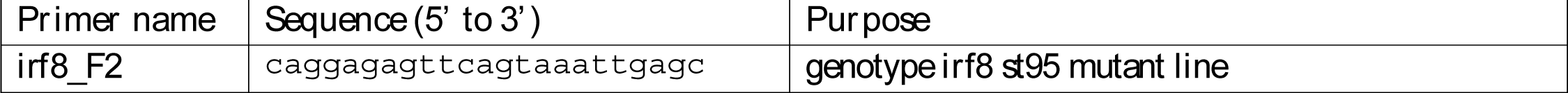
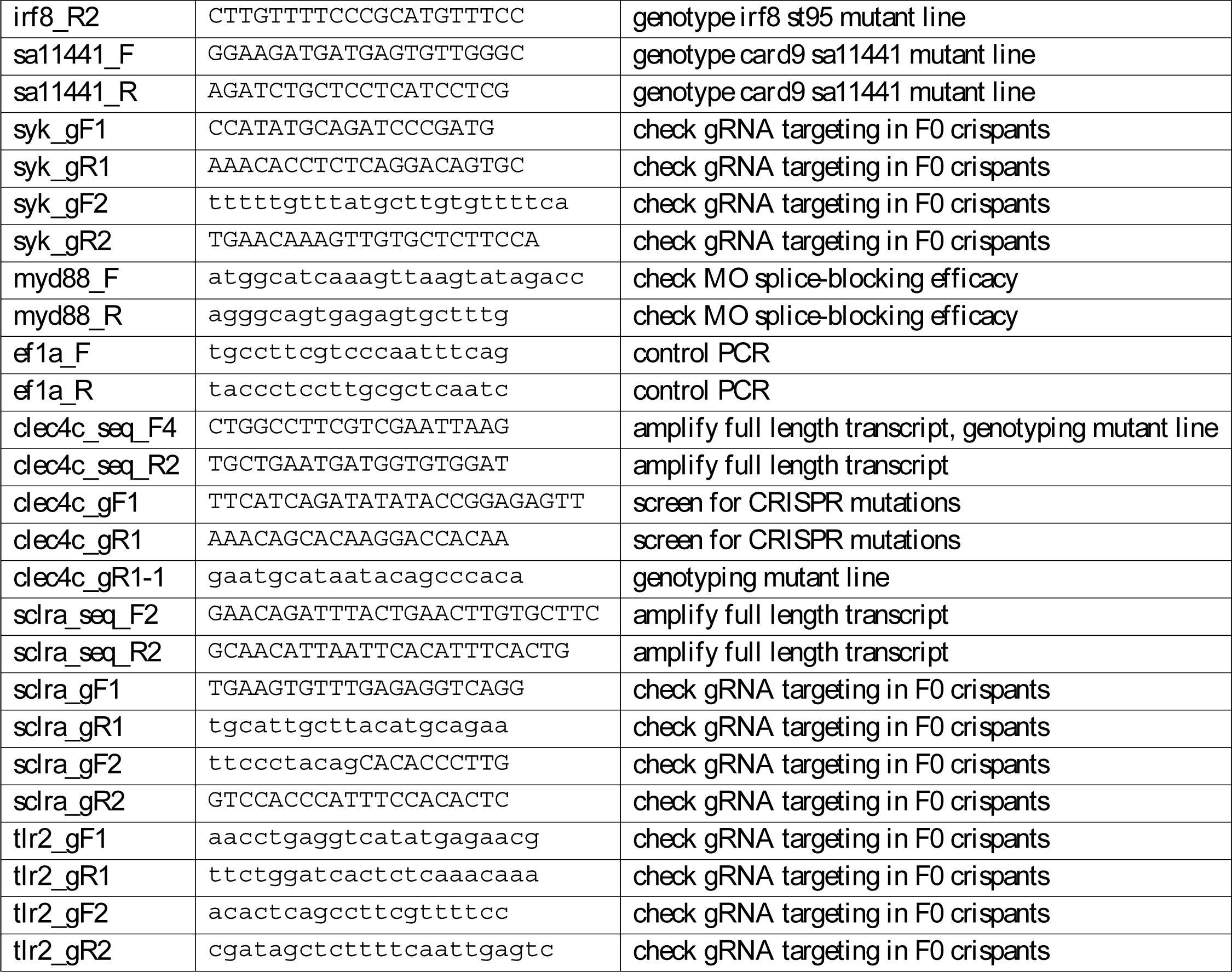
Primers used in this study.

### 2.2 gRNA design and injection

Guide RNAs (gRNAs) were designed with the CHOPCHOP web tool (Labun et al., 2016, Labun et al., 2019, Montague et al., 2014), and were chosen to target sequence encoding required domains of the protein annotated in the human homolog on UniProt (UniProt Consortium, 2023) or in the predicted zebrafish sequence as predicted by Pfam (Mistry et al., 2021) (Table 3, Supp Fig 1A, 2A, 2B, 3). Two gRNAs targeting the *luciferase* coding sequence were also designed as controls. To make gRNAs, DNA oligos were used to generate a double-stranded DNA template containing a T7 promoter sequence (5’-TAATACGACTCACTATAG −3’), the gRNA target sequence without the PAM (Table 3), and the Cas9 scaffold sequence (5’-GTTTTAGAGCTAGAAATAGCAAGTTAAAATAAGGCTAGTCCGTTATCAACTTGAAAAAGTGGCACCGAGTCGGTGCTTTT −3’). Two oligos with overlapping complementarity were annealed, extended with T4 DNA polymerase (NEB), and column purified. In vitro transcription was then done with the HiScribe T7 High Yield RNA Synthesis Kit (NEB), template DNA was removed with DNase I (NEB), and gRNAs were purified with the Monarch RNA Cleanup Kit (NEB), aliquoted, and stored at −80°C. A microinjection setup (BTX, Microject 1000A) supplied with pressure injector, micromanipulator (Narishige), micropipet holder, footswitch, and compressed nitrogen gas was used to inject ∼2 nl of a mix containing 100 ng/µl of each gRNA, 250 µg/ml of Cas9 protein with an NLS resuspended in sterile water (PNA Bio, CP01), and 0.25% phenol red into 1-4 cell stage naturally spawned embryos.

**Table 3.**
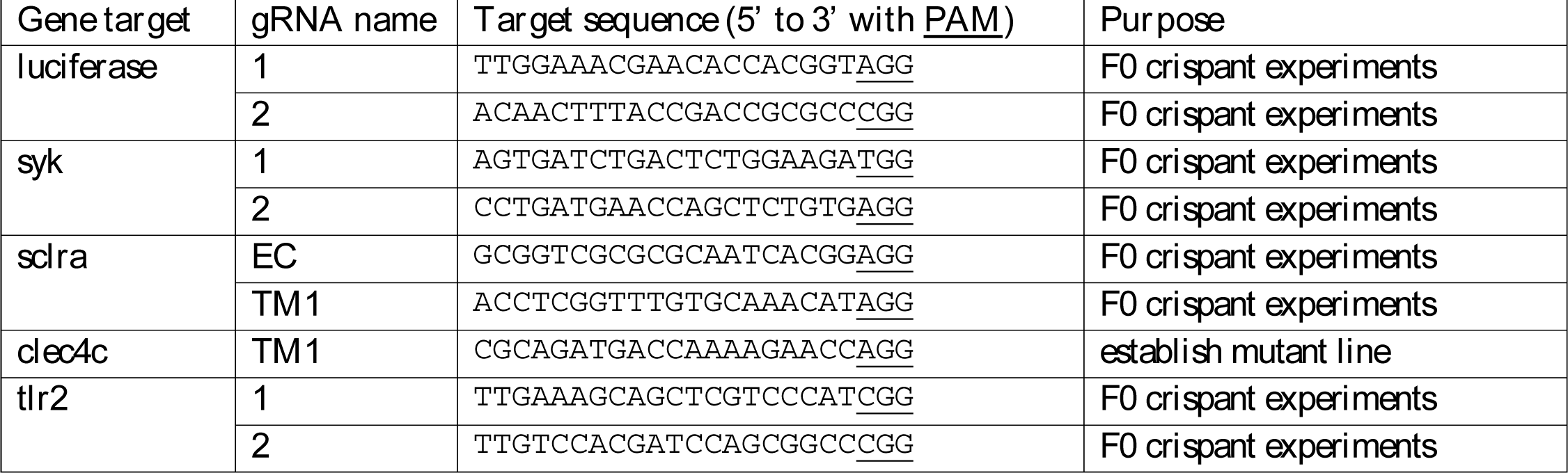
Guide RNA target sites.

For F0 crispant experiments, two guide RNAs targeting the same gene were injected together in the same mix. Primers were designed to amplify ∼100 bp flanking each target site using Primer3 (Untergasser et al., 2012) (Table 2). In every experiment, to confirm successful cutting, which results in DNA repair mechanisms that introduce random small deletions and insertions, PCR was done on gDNA from at least 6 individual larvae using GoTaq Green Master Mix (Promega) (Supp Fig 1A, 2B, 3). PCRs were also performed to detect deletion of the entire DNA region between the two gRNA sites.

### 2.3 Generation of clec4c mutant line

To generate a stable mutant line in *clec4c*, embryos from wild-type adults were injected with a single gRNA (Table 3), and grown to adults. F0 adults were singly outcrossed, gDNA was isolated from eight resulting embryos, and the region around the gRNA site was PCR amplified using GoTaq Green Master Mix (Promega) (primers in Table 2). For F0 adults in which insertions and/or deletions were detected in the germline, embryos from the remaining outcross clutch were grown to adults. F1 adults were then fin clipped, gDNA was isolated, the region around the gRNA site was PCR amplified, cloned into pCR2.1 by TA cloning (Invitrogen), and multiple clones were sequenced to identify specific mutations. A line with a 7 bp deletion (d7) was established (Supp Fig 2A) and outcrossed to the *NF-*κ*B RE::EGFP* reporter (Table 1). This d7 mutation abolishes a BstNI restriction enzyme site and this line was genotyped by PCR with primers clec4c_seq_F4 and clec4c_gR1-1 (Table 2) which amplify 380 bp, followed by a BstNI digest (NEB) (Supp Fig 2A).

### 2.4 Morpholino injection

A *myd88* morpholino (ZFIN MO2: 5’-gttaaacactgaccctgtggatcat −3’) (GeneTools) was previously published and found to be effective at altering splicing of *myd88* mRNA after injection of ∼3 nl at a concentation of 0.33 mM (Bates et al., 2007, Rosowski, Raffa et al., 2018). A standard control MO (GeneTools) was injected at a matching concentration. MO were stored at 4°C in water at 1 mM and diluted for injections with 0.5X CutSmart Buffer (NEB) and 0.1% phenol red. MO mixes were injected into embryos with a microinjection setup (BTX, Microject 1000A) supplied with pressure injector, micromanipulator (Narishige), micropipet holder, footswitch, and compressed nitrogen gas. To confirm splicing changes, RNA was isolated from individual larvae with TRIzol according to the manufacturer’s instructions (Invitrogen) and cDNA was made with iScript RT Supermix (Bio-Rad). The transcript was PCR amplified (primers in Table 2) and gel electrophoresis was done to determine the inclusion of intron 2 (Supp Fig 1C). Primers for *ef1a* were also used as a control (Table 2) (Oehlers et al., 2010).

### 2.5 Cloning and sequencing of putative CLR transcripts

The gene *clec4c* (NCBI Gene ID: 563797, si:dkey-26c10.5 in GRCz11) has two predicted RefSeq transcripts (Supp Fig 2A). To confirm expression of this gene, RNA was isolated from zebrafish larvae using TRIzol according to the manufacturer’s instructions (Invitrogen) and cDNA was made with iScript RT Supermix (Bio-Rad). PCR with Q5 High-Fidelity DNA Polymerase (NEB) and primers clec4c_seq_F4 and clec4c_seq_R2 (Table 2) were used to amplify the shorter predicted transcript (Supp Fig 2C). To sequence this amplified transcript, it was cloned into the pCR Blunt vector with a ZeroBlunt PCR cloning kit (Invitrogen) according to the manufacturer’s instructions. Plasmid DNA from three colonies with inserts were sequenced (Eton Bioscience).

The gene *sclra* (NCBI Gene ID: 564061, si:ch73-86n18.1 in GRCz11, now renamed *cldc1*) has one predicted RefSeq transcript (Supp Fig 2B). To confirm expression of this gene, RT-PCR was performed as above with primers sclra_seq_F2 and sclra_seq_R2 (Table 2) (Supp Fig 2C) and this transcript was cloned as above. Plasmid DNA from eight colonies with inserts were sequenced (Eton Bioscience).

### 2.6 Zymosan injections

Zymosan A, derived from *Saccharomyces cerevisiae* (Sigma-Aldrich), was resuspended in a 1:1 solution of water and DMSO at a concentration of 100 mg/ml. Zymosan was labeled with AlexaFluor633 as previously described for fungal spores (Jhingran et al., 2012, Rosowski, Raffa et al., 2018). Briefly, zymosan particles were resuspended in 0.05 M NaHCO3 and incubated with biotin-XX, SSE (Molecular Probes) at 4°C for ∼4 hours.

Zymosan particles were then washed twice with 100 mM Tris-HCl pH 8.0 to deactivate free-floating biotin and incubated with streptavidin-AlexaFluor633 (Invitrogen) in PBS, washed, resuspended in water/DMSO as for unlabeled particles, and stored at 4°C for up to ∼2 weeks. Zymosan stocks were diluted 1:1 with phenol red to track successful injection and injected into the hindbrain of larvae as previously described (Thrikawala, Savini and Rosowski, 2020). Anesthetized 2 days post fertilization (dpf) larvae were placed on an agarose plate on their lateral side for injection. A microinjection setup (Applied Scientific Instrumentation, MPPI-3) supplied with a pressure injector, micromanipulator (Narishige), micropipet holder, footswitch, back pressure unit, and compressed filtered air was used to inject ∼2 nl of the zymosan suspension into the hindbrain ventricle of each larva. After injection, larvae were rinsed at least twice with E3 to remove tricaine and any free particles.

### 2.7 Microscopy and image analysis

Larvae were screened for fluorescent transgene expression (*NF*κ*B RE::EGFP*, *lyz::BFP*, *mpeg1::mCherry-H2B*, and/or *mfap4::BFP*) on a Zeiss SteREO Discovery.V12 microscope prior to zymosan injection. After zymosan injection, larvae were mounted in glass-bottom dishes in 1% low-melting-point agarose (Fisher BioReagents) and imaged using a Zeiss Cell Observer Spinning Disk confocal microscope on a Axio Observer 7 microscope stand with a confocal scanhead (Yokogawa CSU-X), a Photometrics Evolve 512 EMCCD camera, and a Plan-Apochromat 10x objective (0.3 NA). ZEN software was used to acquire Z-stack images of the hindbrain area every 5 µm. All images were analyzed with ImageJ/Fiji (Schindelin et al., 2012). For timelapse analysis, maximum intensity Z projections of the entire stack were generated and an ROI was drawn to encompass the hindbrain. Mean intensity of EGFP signal was quantified and auto-thresholding with the Otsu algorithm was done to quantify the area of macrophage (*mpeg1::mCherry-H2B*) and neutrophil (*lyz::BFP*) signal. Data was then normalized to the first timepoint. For single timepoint imaging at 8 hours post injection (hpi), maximum intensity Z projections of 20 slices were generated and the mean intensity of EGFP was quantified from an ROI of the hindbrain. All displayed images are processed with bilinear interpolation to increase the pixel density two-fold and some display EGFP intensity with the Fire lookup table (LUT) to better visualize intensity differences.

### 2.8 Statistical analysis

EGFP intensity was analyzed by ANOVA in R version 3.5.2. For each condition, estimated marginal means (emmeans) and SEM were calculated from three independent replicates, each consisting of multiple larvae, and pairwise comparisons were performed with Tukey’s adjustment. Graphs representing data comparing EGFP expression in different genetic backgrounds show all data points, with each data point representing one larvae, color-coded by replicate. P values indicating comparisions to the control-injected condition in the same genetic background are indicated next to emmeans of zymosan-injected conditions. P values of comparisons between genetic backgrounds are indicated by brackets. * = p<0.05, ** = p<0.01, *** = p<0.001, **** = p<0.0001.

## 3. Results

### 3.1 Zebrafish larvae mount an innate immune response to zymosan particles

To begin to characterize the ability of zebrafish to sense and respond to fungal cell wall ligands, and β-glucan in particular, we injected the hindbrain of 2 days post fertilization (dpf) wild-type zebrafish larvae with zymosan particles which primarily consist of β-glucan derived from the cell wall of *Saccharomyces cerevisiae* (Fig 1A). We injected these particles in triple transgenic larvae in which macrophage nuclei (*mpeg1::mCherry*-*H2B*) and neutrophils (*lyz::BFP*) are labeled and which express EGFP when NF-κB transcription factors are activated (*NF*κ*B RE::EGFP*). We then used timelapse microscopy to monitor macrophage and neutrophil recruitment to the area as well as NF-κB activation. Neutrophils arrive to the injection site first but are rapidly followed by macrophages (Fig 1B,C, Supp Video 1). Neutrophils reach their peak recruitment around 400-500 minutes post injection and then begin to decrease. Macrophages, on the other hand, have sustained and higher recruitment, peaking at ∼600 minutes post injection and then are maintained in the hindbrain (Fig 1B,C, Supp Video 1). In addition to cell recruitment, we also observed EGFP expression, indicating NF-κB activation (Fig 1B,C, Supp Video 1). This EGFP expression takes longer to be detectable, however this is partially due to the time it takes for mRNA transcription, translation, and protein folding, and we cannot make any conclusions about the timing of NF-κB activation from this experiment. These data demonstrate that zebrafish larvae can mount an innate immune response to zymosan and provide further evidence that zebrafish have a receptor for β-glucan.

**Figure 1.**
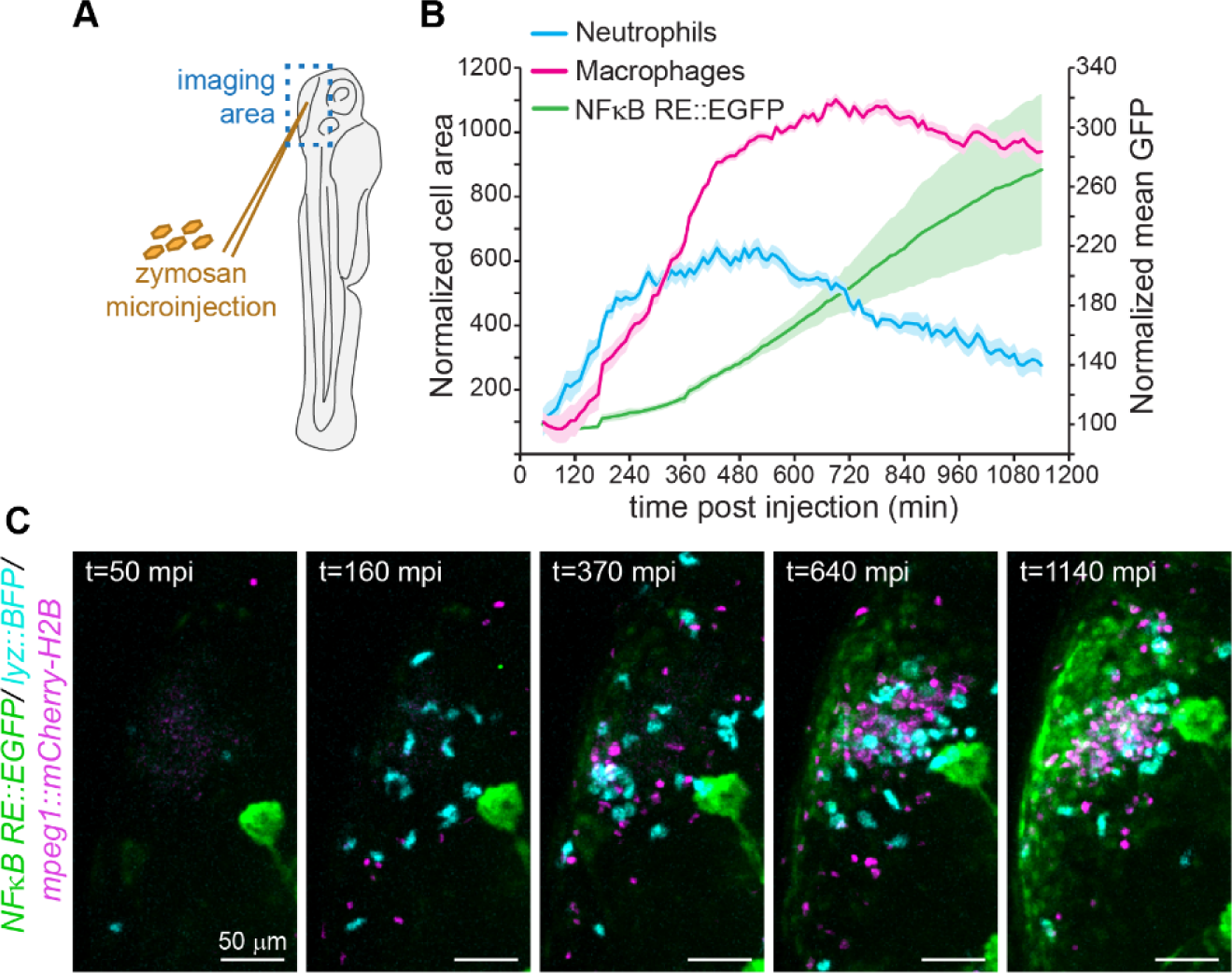
Zebrafish larvae mount an innate immune response to zymosan particles. **A.** Schematic of injection of zymosan particles into hindbrain of 2 dpf zebrafish larvae and subsequent live imaging. **B.** Transgenic larvae with labeled neutrophils (*lyz::BFP*), labeled macrophage nuclei (*mpeg1::mCherry-H2B*), and that express EGFP when NF-κB transcription factors are activated (*NF*κ*B RE::EGFP*) were injected with zymosan particles and live imaged by timelapse microscopy every 10 min. Neutrophil, macrophage, and EGFP signal is plotted, normalized to the initial time point at 50 mpi. Lines represent average across five larvae with shading representing standard deviation. **C.** Example images from one larvae from timelapse microscopy.

### 3.2 Macrophages and neutrophils mediate the response to zymosan

We next wondered what the role of macrophages and neutrophils was in this response and in particular whether both cell types can phagocytose zymosan particles and if NF-κB is activated within both cell types. To investigate these questions, we turned again to timelapse microscopy after injection of zymosan labeled with an AlexaFluor molecule (AF633). First, we used larvae that have a fluorescent label in the cytosol of macrophages (*mfap4::BFP*) as well as the NF-κB reporter (Fig 2A, Supp Video 2). From this timelapse imaging we observed that macrophages can phagocytose zymosan particles (Fig 2Ai) and we also observed NF-κB activation inside of macrophages (Fig 2Aii). Similarly, in an experiment with larvae that have fluorescently-labeled neutrophils (*lyz::BFP*) as well as the NF-κB reporter (Fig 2B, Supp Video 3), we observe neutrophils phagocytosing zymosan particles (Fig 2Bi,ii) and expressing EGFP downstream of NF-κB activation (Fig 2Biii). These results demonstrate that both macrophages and neutrophils can respond to zymosan.

**Figure 2.**
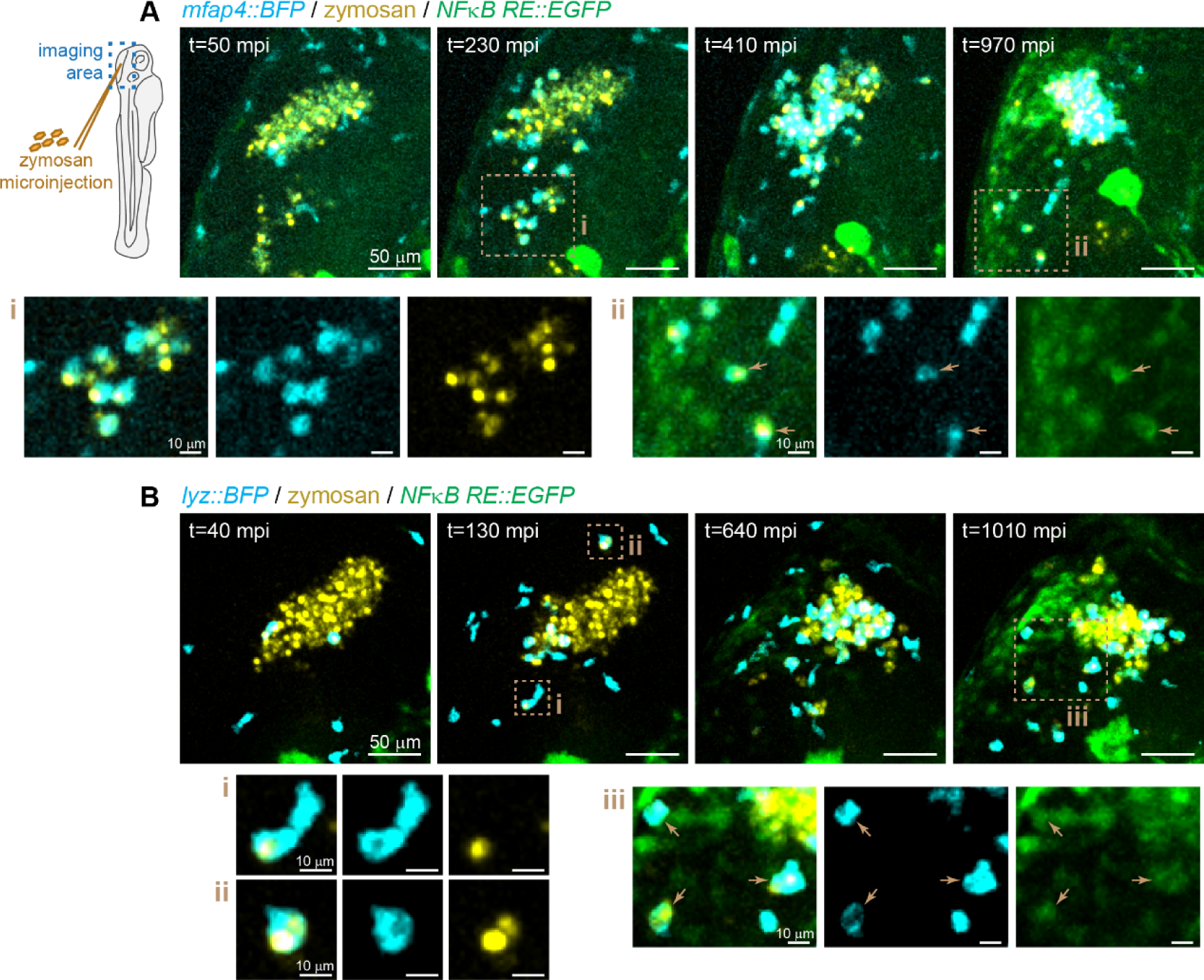
Macrophages and neutrophils phagocytose zymosan and activate NF-κB. **A.** Transgenic larvae with labeled macrophages (*mfap4::BFP*), and that express EGFP when NF-κB transcription factors are activated (*NF*κ*B RE::EGFP*) were injected with AF633-labeled zymosan particles at 2 dpf and live imaged by timelapse microscopy every 10 min. Images from one representative larvae are shown. **B.** Transgenic larvae with labeled neutrophils (*lyz::BFP*), and that express EGFP when NF-κB transcription factors are activated (*NF*κ*B RE::EGFP*) were injected with AF633-labeled zymosan particles at 2 dpf and live imaged by timelapse microscopy every 10 min. Images from one representative larvae are shown.

However, it was still unclear if macrophages and neutrophils were sensing and responding to the zymosan directly or if other cells, like epithelial cells, sense the zymosan, activate NF-κB, and secrete chemokines to recruit macrophages and neutrophils. Indeed, from our timelapse imaging there appears to be significant NF-κB signal that does not overlap with either macrophages or neutrophils (Fig 1C). We therefore wanted to test if macrophages and/or neutrophils are required for NF-κB activation in response to zymosan or if other cells, like epithelial cells, can sense and activate NF-κB independently. To that end, we combined a zebrafish neutrophil defective model (*mpx::mCherry-2A-rac2D57N*; hereafter referred to as *rac2D57N*)(Deng et al., 2011) with a macrophage-deficient model (*irf8*^-/-^)(Shiau et al., 2015) alongside the NF-κB reporter. By combining these models we could monitor NF-κB activation in larvae without macrophages, without functioning neutrophils, or without either functional cell type, all from the same cross (Fig 3A). We injected zymosan or vehicle control into these larvae and imaged the hindbrain for EGFP expression at 8 hours post injection (hpi). In all control vehicle-injected larvae, little EGFP expression was observed in the hindbrain (Fig 3B). In wild-type (*irf8*^+/+^; no transgene) larvae, significant EGFP expression occurred (Fig 3B,C), in line with results from our timelapse experiment (Fig 1B). In larvae lacking macrophages but having functional neutrophils (*irf8*^-/-^; no transgene), statistically significantly lower NF-κB activated EGFP expression occurred (Fig 3B,C), indicating that macrophages play an important role in responding to zymosan. Surprisingly, in larvae that lack functional neutrophils but have macrophages (*irf8*^+/+^; *rac2D57N*), there was no decrease in NF-κB activated EGFP expression (Fig 3B,C), demonstrating that macrophages are sufficient and neutrophils are not required for NF-κB activation in response to zymosan in larval zebrafish. However, in larvae that do not have macrophages, loss of functional neutrophils (*irf8*^+/+^; *rac2D57N*) leads to little NF-κB activation above baseline (Fig 3B,C), demonstrating that in the absence of macrophages, neutrophils play an important role in responding to this PAMP. Altogether, these data demonstrate that innate immune cells are absolutely required for NF-κB activation in response to zymosan. While neutrophils can play a role in this response, macrophages on their own are sufficient to sense this PAMP and activate NF-κB at the injection site.

**Figure 3.**
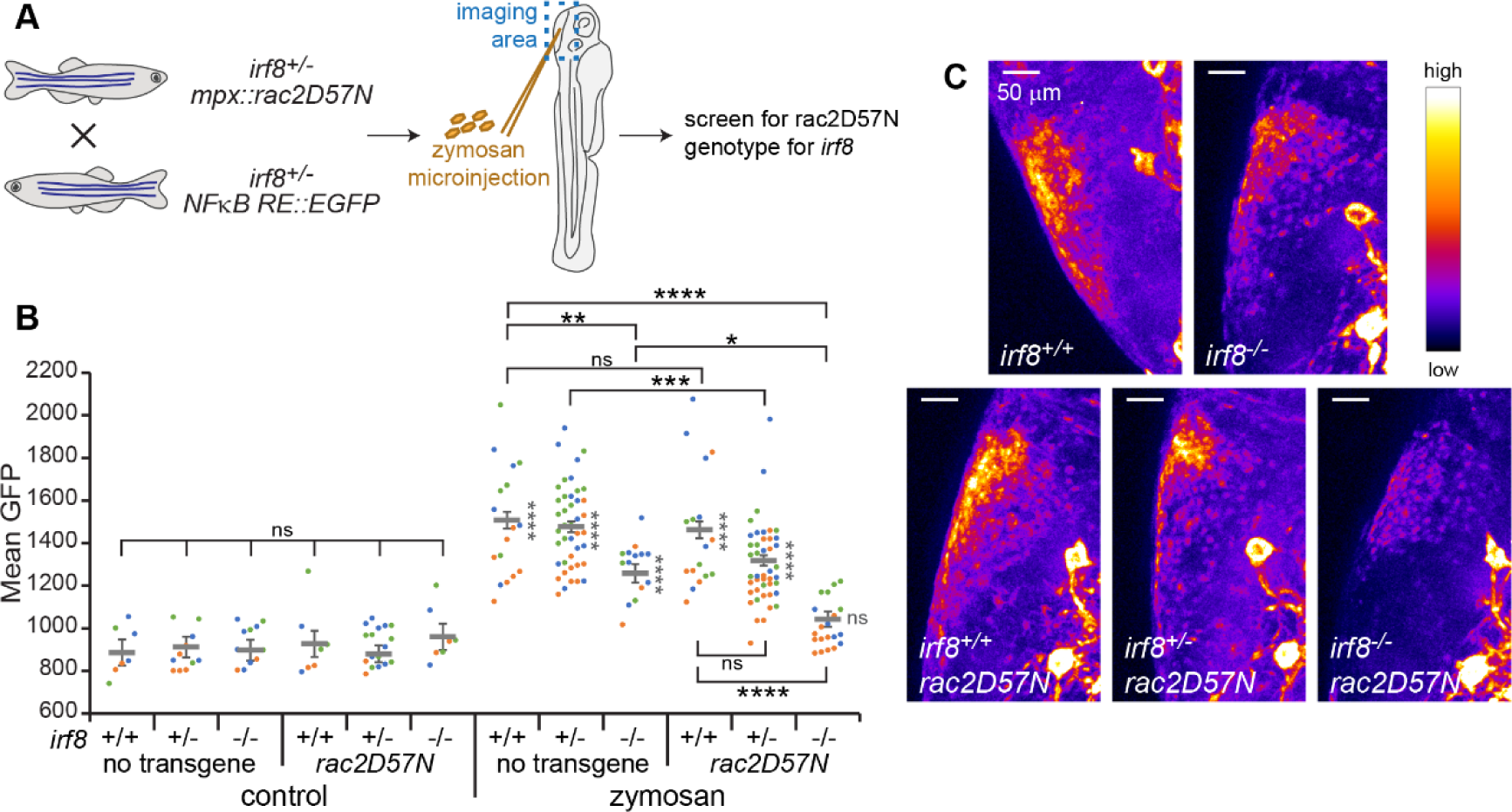
Macrophages and neutrophils are required for NF-κB activation in response to zymosan. **A.** Experimental setup. Adult zebrafish heterozygous for the *irf8* transcription factor required for macrophage development and carrying either the *mpx::rac2D57N* transgene which makes neutrophils defective or the *NF*κ*B RE::EGFP* transgene which marks NF-κB activation were crossed. Resulting larvae were injected with zymosan particles at 2 dpf and live imaged at 8 hpi. Larvae were screened for mCherry fluorescence to determine the presence of the *rac2D57N* transgene and genotyped at the *irf8* locus from gDNA after imaging. **B.** EGFP fluorescence in the hindbrain was quantified from images. Data represent 3 pooled replicates. Each symbol represents one larva, color-coded by replicate. Lines represent emmeans ± SEM. **C.** Example images showing EGFP intensity with the Fire LUT.

### 3.3 Conserved CLR signaling pathways are used in larval zebrafish to sense zymosan

As the receptors that bind to PAMPs in zymosan are still unknown in zebrafish, we first aimed to determine if canonical signaling pathways downstream of PRRs contribute to the response to zymosan by interfering with genes encoding adaptor proteins downstream of PRRs (Fig 4A). We first used a CRISPR/Cas9 system to generate crispants in *syk*, a kinase downstream of C-type lectin receptor (CLR) activation that is upstream of NF-κB activation (Supp Fig 1A). We injected zymosan into *syk* crispant larvae or larvae derived from embryos injected with a gRNA targeting *luciferase* as a control, all in the NF-κB reporter background, and imaged the hindbrain at 8 hpi. We found that in larvae with *syk* targeted, NF-κB activation in response to zymosan injection was statistically significantly lower than in control *luc* gRNA larvae (Fig 4B, C). However, there is still significant EGFP expression compared to vehicle injected controls (Fig 4B). Another conserved signaling protein downstream of CLR activation is Card9, and we also crossed a *card9* mutant line from the Sanger mutation project to the NF-κB reporter line (Supp Fig 1B). In *card9*^-/-^ larvae, NF-κB activation in response to zymosan is also significantly lower than in *card9*^+/-^ siblings, while these larvae still have significant EGFP expression compared to vehicle injected controls (Fig 4D, E). These experiments indicate that signaling through Card9 and Syk promotes NF-κB activation in response to zymosan, suggesting that zebrafish have a CLR that can sense zymosan. However, there are likely other pathways that can also sense zymosan.

**Figure 4.**
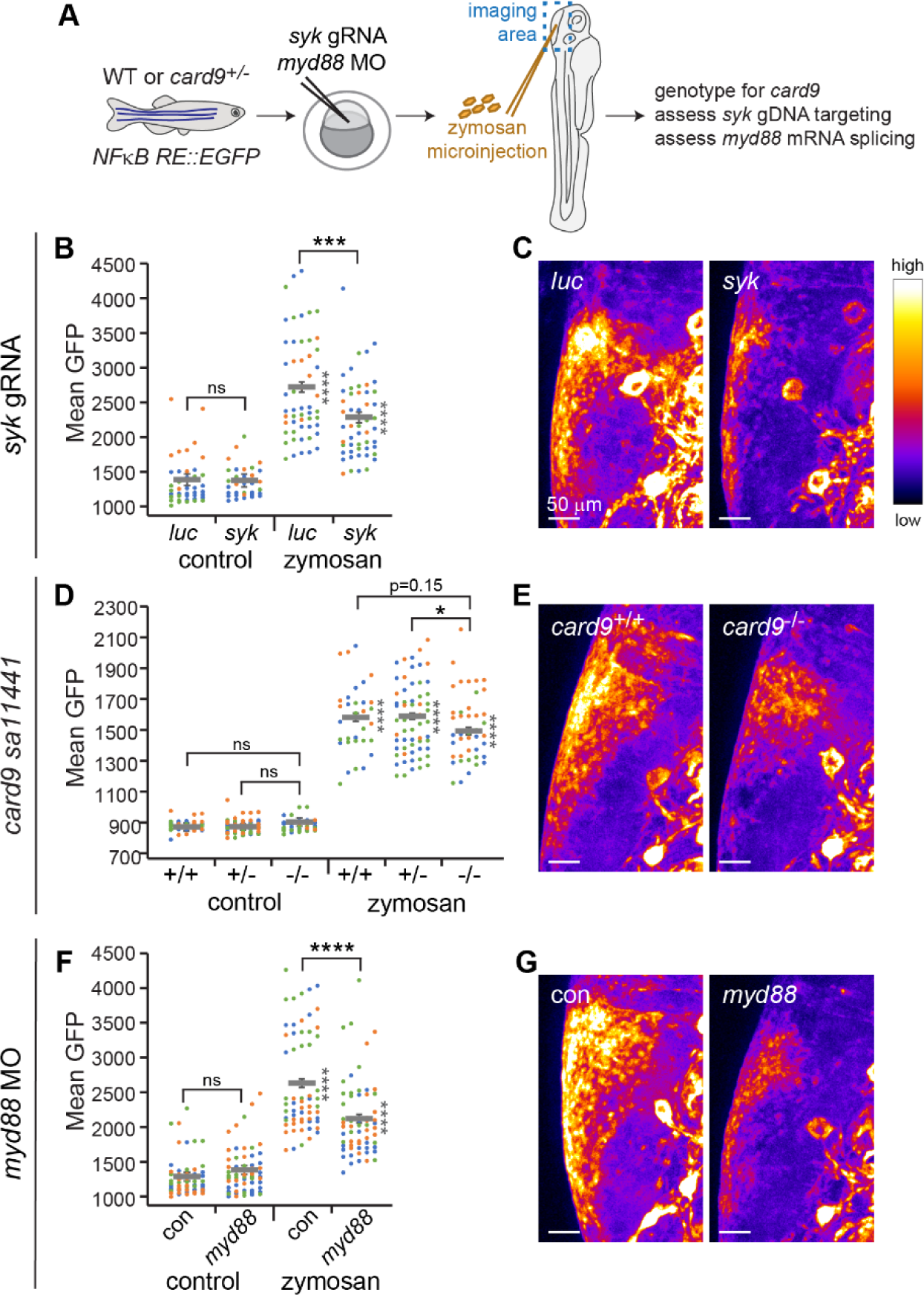
Zymosan activates conserved CLR and TLR signaling pathways. **A.** Experimental setup. In the *NF*κ*B RE::EGFP* transgenic line, genes encoding adaptor proteins were genetically manipulated by stable mutation in adults (*card9*), injection of gRNAs and Cas9 into embryos (*syk*), or injection of morpholino into embryos (*myd88*). At 2 dpf, larvae were injected with zymosan particles and live imaged at 8 hpi. Larvae were genotyped at the *card9* locus, *syk* gDNA targeting was confirmed, or *myd88* MO efficiency was monitored. **B, C.** *syk* targeting by CRISPR/Cas9. gRNAs against *luciferase* were used as a control. (B) EGFP fluorescence in the hindbrain was quantified from images. Data represent 3 pooled replicates. Each symbol represents one larva, color-coded by replicate. Lines represent emmeans ± SEM. (C) Example images showing EGFP intensity with the Fire LUT. **D, E.** *card9* mutation. (D) EGFP fluorescence in the hindbrain was quantified from images. Data represent 3 pooled replicates. Each symbol represents one larva, color-coded by replicate. Lines represent emmeans ± SEM. (E) Example images showing EGFP intensity with the Fire LUT. **F, G.** *myd88* morpholino. A standard control morpholino was used as a control. (F) EGFP fluorescence in the hindbrain was quantified from images. Data represent 3 pooled replicates. Each symbol represents one larva, color-coded by replicate. Lines represent emmeans ± SEM. (G) Example images showing EGFP intensity with the Fire LUT.

### 3.4 TLR signaling pathways are also used in larval zebrafish to sense zymosan

As our data suggests that other non-CLR PRRs can also sense and initiate responses to zymosan in larval zebrafish, we next decided to test the requirement of a signaling adaptor downstream of TLR activation, Myd88. We used an established morpholino (MO) to interfere with *myd88* mRNA splicing and prevent correct translation (Supp Fig 1C). In larvae with *myd88* knocked down, NF-κB activation in response to zymosan was significantly lower than control MO larvae (Fig 4F, G), demonstrating that TLR signaling can also be used by larval zebrafish to sense zymosan. However, again, activation in *myd88* larvae was still above vehicle control-injected baseline. Altogether, we find that zebrafish larvae can use both CLR and TLR signaling to activate NF-κB in response to zymosan and thus likely have multiple receptors that can bind to the PAMPs in zymosan.

### 3.5 Putative C-type lectin receptors are required for sensing of zymosan

Recently, Petit et al identified two putative CLR genes on zebrafish chromosome 16 in a region that shares synteny with the genomic locations of three CLRs in human and mouse genomes, Dectin-1, Dectin-2, and Mincle (Petit et al., 2019). These genes were named *clec4c* (NCBI Gene ID: 563797, si:dkey-26c10.5 in GRCz11) and *sclra* (NCBI Gene ID: 564061, si:ch73-86n18.1 in GRCz11, now renamed *cldc1*) (Supp Fig 2A,B). We first confirmed that transcripts of both of these genes are expressed in larval zebrafish (Supp Fig 2C). *clec4c* has two transcripts predicted by RefSeq, but were only able to amplify the shorter transcript, and sequencing of three clones of this transcript confirmed that it perfectly matched the RefSeq prediction. Seven of the eight cloned transcripts of *sclra* also had identical sequence to the single RefSeq prediction for this gene, with one transcript missing 61 nucleotides at the beginning of exon 3.

To test the hypothesis that one or both of these putative CLRs promotes sensing of zymosan in zebrafish, we again used CRISPR/Cas9 to target these genes. We first generated a stable mutant line in *clec4c* which contains a 7 bp deletion in the predicted transmembrane domain which leads to a frameshift mutation and premature stop codon within this domain (Supp Fig 2A), and crossed this mutant line to the transgenic NF-κB reporter. We then generated crispants in *sclra* (Supp Fig 2B) in progeny from an in-cross of *clec4c* mutants with the NF-κB reporter to generate larvae that are *clec4c* mutants, *sclra* crispants, or both, all from one clutch (Fig 5A). After injection with zymosan, *clec4c*^-/-^ larvae had lower NF-κB-activated GFP expression than wild-type or heterozygous siblings, although this difference was not quite statistically significant (Fig 5B, C). Similarly, *sclra* crispant larvae in a wild-type background had statistically significantly lower NF-κB-activated GFP expression after zymosan injection than *luciferase*-targeted controls (Fig 5B, C). Interestingly, larvae that were both homozygous mutants in *clec4c* and *sclra*-targeted had similar levels of GFP expression to larvae with only one of these genes targeted (Fig 5B, C). These data suggest that both *clec4c* and *sclra* are required for full NF-κB activation in response to zymosan and that these genes likely work in the same pathway to sense fungal PAMPs.

**Figure 5.**
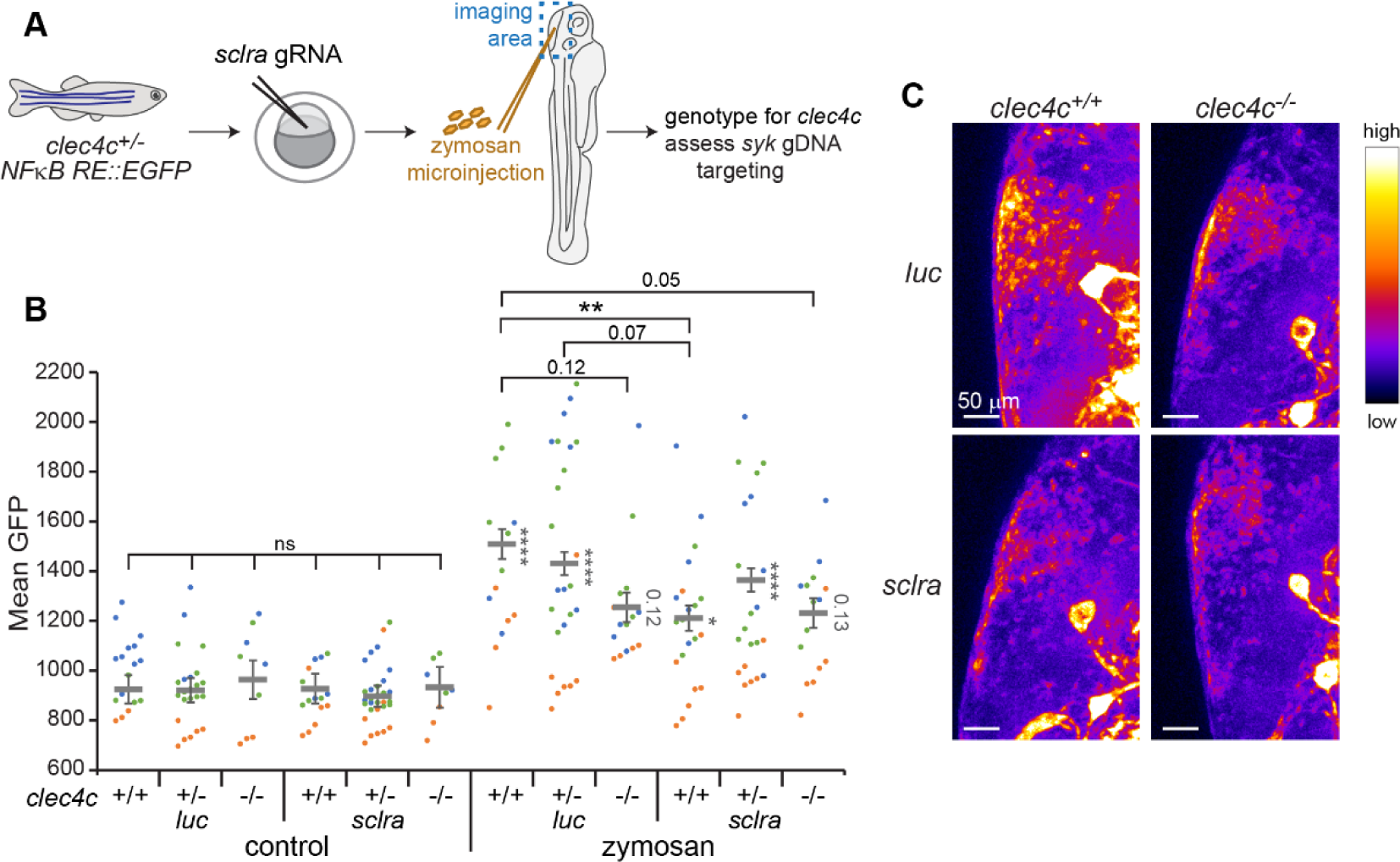
Putative Dectin-1 homologs promote the response to zymosan. **A.** Experimental setup. Adult zebrafish heterozygous for *clec4c* d7 mutation and carrying the *NF*κ*B RE::EGFP* transgene which marks NF-κB activation were crossed. Embryos were injected with Cas9 and gRNAs targeting *sclra* or *luciferase* as a control. At 2 dpf, larvae were injected with zymosan particles and live imaged at 8 hpi. Larvae were genotyped at the *clec4c* locus, and *sclra* gDNA targeting was confirmed. **B.** EGFP fluorescence in the hindbrain was quantified from images. Data represent 3 pooled replicates. Each symbol represents one larva, color-coded by replicate. Lines represent emmeans ± SEM. **C.** Example images showing EGFP intensity with the Fire LUT.

### 3.6 Tlr2 can promote zymosan sensing in the absence of Card9 signaling

As our previous results demonstrated that Myd88 and TLR signaling are also required for maximum NF-κB activation in response to zymosan, we also decided to test whether zebrafish Tlr2 can promote sensing of this fungal particle. Human TLR2 can directly bind zymosan (Sato, M. et al., 2003) and murine Tlr2 is recruited to macrophage phagosomes containing zymosan (Underhill et al., 1999), but whether zebrafish Tlr2 has these roles is unknown. We therefore first used CRISPR/Cas9 to generate *tlr2* crispants (Fig 6A, Supp Fig 3). When we injected *tlr2* crispant larvae with zymosan, we found no difference in NF-κB activated GFP expression compared to *luciferase* gRNA-injected control larvae (Fig 6B), demonstrating that *tlr2* is not required for zebrafish to sense zymosan. We speculated that this might be due to the fact that these larvae still have CLR signaling which would compensate for any effect of the loss of Tlr2. Therefore, to test whether Tlr2 can promote NF-κB activation if CLR signaling is inhibited, we generated *tlr2* crispants in an in-cross of the *card9* mutant line (Fig 6A). In this experiment, again we saw no effect of *tlr2* targeting alone (Fig 5C, D). However, in *card9^+/-^* and *card9*^-/-^ larvae, *tlr2* targeting did decrease NF-κB mediated GFP expression in response to zymosan (Fig 5C, D), suggesting that Tlr2 in zebrafish does play a minor role in zymosan-activated NF-κB-mediated gene expression which is overshadowed by the effect of CLRs.

**Figure 6.**
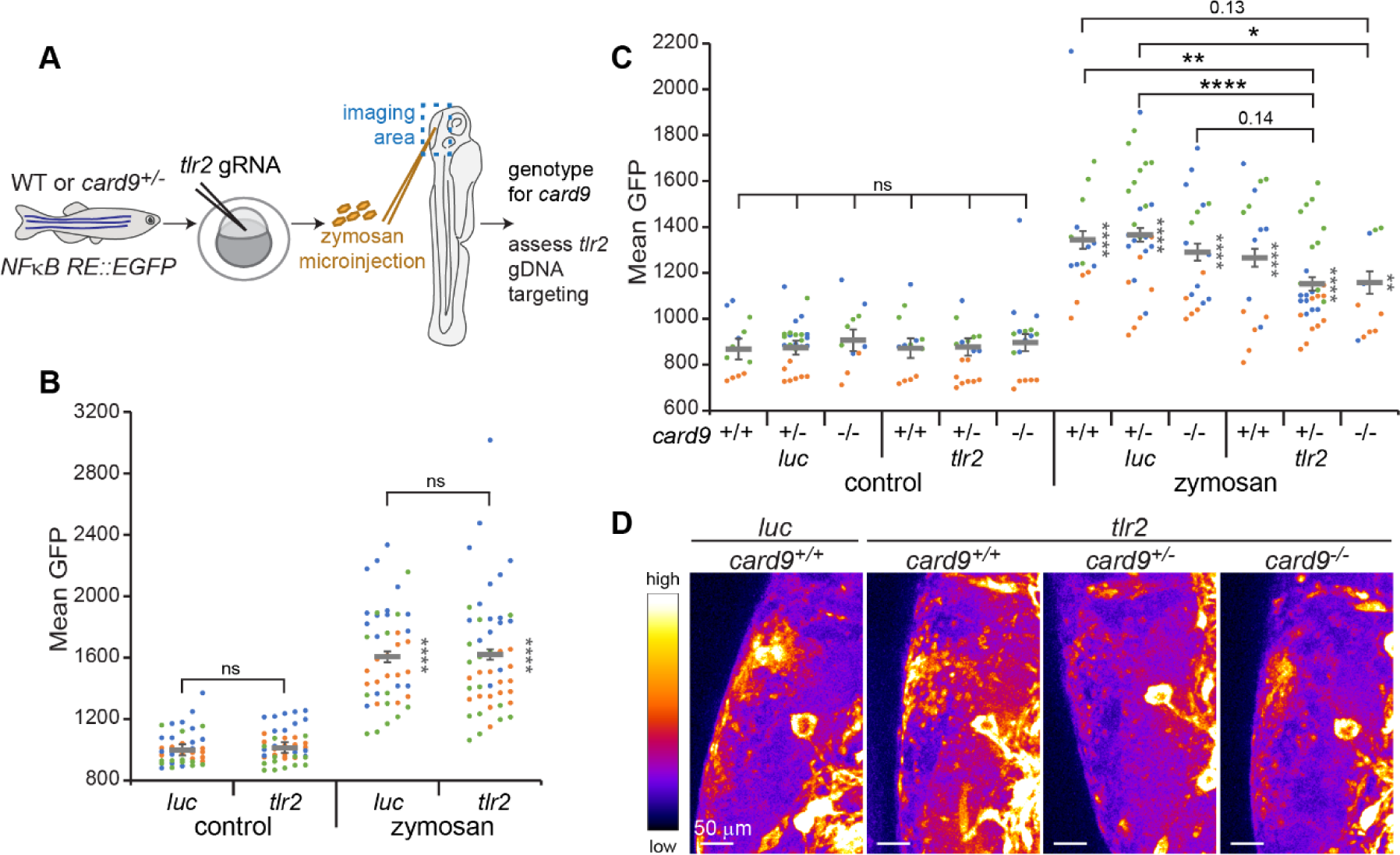
Tlr2 is partially required for the response to zymosan when CLR signaling is inhibited. **A.** Experimental setup. Wild-type adult zebrafish or adults heterozygous for *card9* sa11441 mutation also carrying the *NF*κ*B RE::EGFP* transgene which marks NF-κB activation were crossed. Embryos were injected with Cas9 and gRNAs targeting *tlr2* or *luciferase* as a control. At 2 dpf, larvae were injected with zymosan particles and live imaged at 8 hpi. Larvae were genotyped at the *card9* locus, and *tlr2* gDNA targeting was confirmed. **B.** EGFP fluorescence in the hindbrain was quantified from images of *tlr2* crispant larvae in the wild-type background. Data represent 3 pooled replicates. Each symbol represents one larva, color-coded by replicate. Lines represent emmeans ± SEM. **C.** EGFP fluorescence in the hindbrain was quantified from images of *tlr2* crispant larvae in the *card9* mutant background. Data represent 3 pooled replicates. Each symbol represents one larva, color-coded by replicate. Lines represent emmeans ± SEM. **D.** Example images showing EGFP intensity with the Fire LUT.

## 4. Discussion

Pattern recognition receptors are essential for the initiation of innate immune responses to pathogens. Different receptors families exist, including C-type lectin receptors (CLRs) and Toll-like receptors (TLRs), with conserved domains that allow for identification of members of these families across the evolutionary tree of life (Kang and Lee, 2011, Li, D. and Wu, 2021, Li, Y. et al., 2017, Nie et al., 2018). However, whether receptors in different species recognize the same ligands is unclear (Li, Y. et al., 2017, Nie et al., 2018), posing a problem for modeling infectious diseases in non-human hosts, including zebrafish.

In this study, we investigated the pathways that zebrafish larvae use to sense fungal PAMPs present in zymosan, a particle derived from the cell wall of *Saccharomyces cerevisiae* that primarily contains β-glucan. We report that larvae mount a robust innate immune response to zymosan: macrophages and neutrophils migrate to the site of PAMP injection, both cell types can phagocytose zymosan particles, and NF-κB is activated both in these phagocytes and in the surrounding tissue. Because our experiments utilized an EGFP reporter of NF-κB activation, it takes several hours to observe NF-κB activation but this likely due to the time it takes to transcribe, translate, and fold EGFP rather than a true delay in activation of this signaling pathway and transcription factor. While it was previously found that in larval zebrafish β-glucan can drive increased resistance to bacterial or viral infections (Darroch et al., 2022, Liang et al., 2022) and that zymosan particles or β-glucan-coated beads can be shuttled between phagocytes (Pazhakh et al., 2019), this the first characterization of the innate immune response to these PAMPs in larval zebrafish. The receptors and signaling pathways that zebrafish use to sense fungal PAMPs have been unclear. We find that both CLR and TLR signaling pathways, through the conserved signaling adaptors Card9/Syk and Myd88, respectively, promote NF-κB activation in response to zymosan.

In humans and mice, β-glucan is sensed primarily by the CLR Dectin-1. Searching the zebrafish genome for a predicted gene with significant sequence homology to Dectin-1 has not presented any Dectin-1 homolog candidates. However, Petit et al searched for conserved synteny of region of the genome containing *Dectin-1* in mammals with in the zebrafish genome and identified two putative CLR-encoding genes (Petit et al., 2019). We directly tested the requirement for these two genes, *clec4c* (si:dkey-26c10.5 in GRCz11) and *sclra* (si:ch73-86n18.1 in GRCz11, now renamed *cldc1*), in NF-κB activation in response to zymosan in larval zebrafish. We find that both a stable mutant in *clec4c* and F0 crispants in *sclra* have reduced NF-κB activation in response to zymosan, demonstrating that both of these receptors play a role in responding to fungal PAMPs in zebrafish.

Whether *clec4c* and/or *sclra* are true homologs of *Dectin-1* is still unknown. Zymosan primarily consists of β-glucan and was originally used to identify Dectin-1 in the mouse genome (Brown, G. D. and Gordon, 2001), and Dectin-1 is the receptor in mouse primary macrophages that is primarily responsible for zymosan recognition (Brown, Gordon D. et al., 2002). However, in humans and mice, other cell-surface receptors exist that bind to β-glucan, including Langerin (CD207) (de Jong, Marein A W P et al., 2010), low-affinity IgE receptors CD23 and EphA2 (Guo et al., 2018, Swidergall et al., 2018), Complement receptor 3 (CR3, CD11b/CD18, Mac-1) (Ross et al., 1987), scavenger receptors (Jozefowski et al., 2012), and lactosylceramide (Evans et al., 2005). Transfection of β-glucan into the cytosol of cells can also activate the NLRP3 inflammasome (Briard et al., 2019). Zymosan also contains a low level of mannans (Di Carlo and Fiore, 1958), and some zymosan recognition can be blocked by exogenous mannan (Taylor, Philip R. et al., 2002), thus it is also possible that Clec4c and/or Sclra are recognizing these sugars instead of β-glucan. Mannans are recognized by several PRRs, including Dectin-2 (Feinberg et al., 2017, McGreal et al., 2006, Reedy et al., 2023, Sato, K. et al., 2006). *Dectin-2* is located in the same identified syntenic region of the genome as *Dectin-1* and it is possible that *clec4c* and/or *sclra* is a homolog for *Dectin-2*. The other CLR coded for in this syntenic region, Mincle, recognizes many different ligands but the specific ligands bound that are present in major human fungal pathogens are not known (Hatinguais et al., 2020).

Interestingly, we find that mutation of *clec4c* or *sclra* individually and targeting of both genes together results in identical phenotypes with respect to NF-κB activation in response to zymosan, suggesting that these receptors act in the same pathway to sense the fungal PAMPs present in these particles. CLRs can associate as heterodimers. For example, Macrophage C-type Lectin (MCL, Clec4d, Dectin-3) requires Mincle for expression at the surface of mouse myeloid cells (Kerscher et al., 2016) and Dectin-2 and MCL can also form heterodimers to recognize α-mannans (Zhu et al., 2013). One hypothesis therefore is that Clec4c and Sclra form a heterodimer to sense zymosan, such that mutation of only one of these genes abolishes the function of the heterodimer.

We find that TLR signaling also promotes NF-κB activation in response to zymosan. Mammalian TLR2 recognizes mannosylated PAMPs (Oliveira-Nascimento et al., 2012) and can recognize and bind zymosan (Sato, M. et al., 2003, Underhill et al., 1999), and we similarly identify a role for zebrafish Tlr2 in this response. While TLR2 is not required for the uptake of zymosan by mouse bone marrow derived macrophages, TLR2 and Dectin-1 expression synergistically increase NF-κB activation in HEK293 cells in response to zymosan stimulation (Gantner et al., 2003). We find that in otherwise wild-type larval zebrafish, mutation of *tlr2* does not have an impact on NF-κB activation in response to zymosan. However, in *card9*^-/-^ larvae, in which CLR signaling is inhibited, targeting of *tlr2* does decrease NF-κB activation. These results suggest that Tlr2 in zebrafish plays a secondary role in sensing of fungal PAMPs that is not required for innate immune activation when CLR signaling is intact.

We find that macrophages and neutrophils are required for recognition of zymosan in larval zebrafish. In *irf8* mutant larvae, which lack macrophages, NF-κB reporter line activation is significantly lower than in *irf8*^+/+^ siblings in response to zymosan injection. However, in larvae that have dysfunctional neutrophils (*mpx::rac2D57N*), we do not observe a decrease in NF-κB activation, demonstrating that while macrophages are required for a full response to zymosan, neutrophils on their own are not. However, when macrophages are absent, the loss of neutrophil function brings NF-κB activation in response to zymosan down to baseline levels, demonstrating that while macrophages alone are sufficient to respond to zymosan, neutrophils also play a secondary role in NF-κB activation in response to these PAMPs. It should be noted that Irf8 also has transcriptional roles outside of its requirement in macrophage development and can co-regulate many targets of NF-κB signaling (Liu and Ma, 2006, Simon et al., 2015, Yan et al., 2020, Yan et al., 2022). This may explain why we see a decrease in NF-κB activation in response to zymosan in *irf8* heterozygotes compared to wild-type siblings in the *racD57N* neutrophil-defective background.

The requirement for macrophages and neutrophils in NF-κB activation in response to zymosan is consistent with the idea that the receptor(s) that mediate this recognition are expressed specifically on myeloid cells. Both Dectin-1 and Dectin-2 are predominantly expressed on myeloid cells, however, Dectin-2 in mouse and human neutrophils is only expressed after stimulation with pro-inflammatory cytokines (Taylor, Patricia R. et al., 2014, Taylor, Philip R. et al., 2002). However, in our imaging experiments, GFP signal indicating NF-κB activation can clearly be seen in cells that do not have either macrophages or neutrophil marker expression and that appear to be epithelial cells. In fact, the epithelium is thought to play a role in orchestrating immunity in response to fungi (Heung et al., 2023, Jhingran et al., 2015) and human lung epithelial cells can express Dectin-1 (Heyl et al., 2014, Sun et al., 2012). However, our results indicate that any NF-κB activation in epithelial cells in these experiments must be largely due to secreted signals from phagocytes, rather than to direct sensing of the PAMPs present in zymosan.

We find that in larval zebrafish both macrophages and neutrophils respond to and phagocytose zymosan, but it is still unclear whether the same receptors required for NF-κB activation are required for phagocyte chemotaxis to fungal PAMPs and uptake of zymosan. Additionally, the roles of Clec4c, Sclra, and Tlr2 in resistance to fungal pathogens in larval zebrafish remain to be tested and should be the subject of future studies. Dectin-1 deficient mice are more susceptible to *A. fumigatus* infection (Werner et al., 2009), while the requirement for Dectin-1 against *C. albicans* infection depends on the strain and the mode of infection (Marakalala et al., 2013, Saijo et al., 2007, Taylor, Philip R. et al., 2007, Vautier et al., 2012). Further characterization of these PRRs in larval zebrafish will better define their homology to mammalian CLRs and their role in immunity.

## Acknowledgements

We thank members of the Rosowski Lab for helpful discussions and for assistance with zebrafish care. We thank the Clemson University Aquatic Animal Research Laboratory for providing the microinjection setup used for single-cell embryo injections. We thank the Zebrafish International Resource Center for providing the *card9* sa11441 mutant line. We thank John Rawls for sharing the NF-κB reporter transgenic line.

---

**Supplementary Figure 1.**
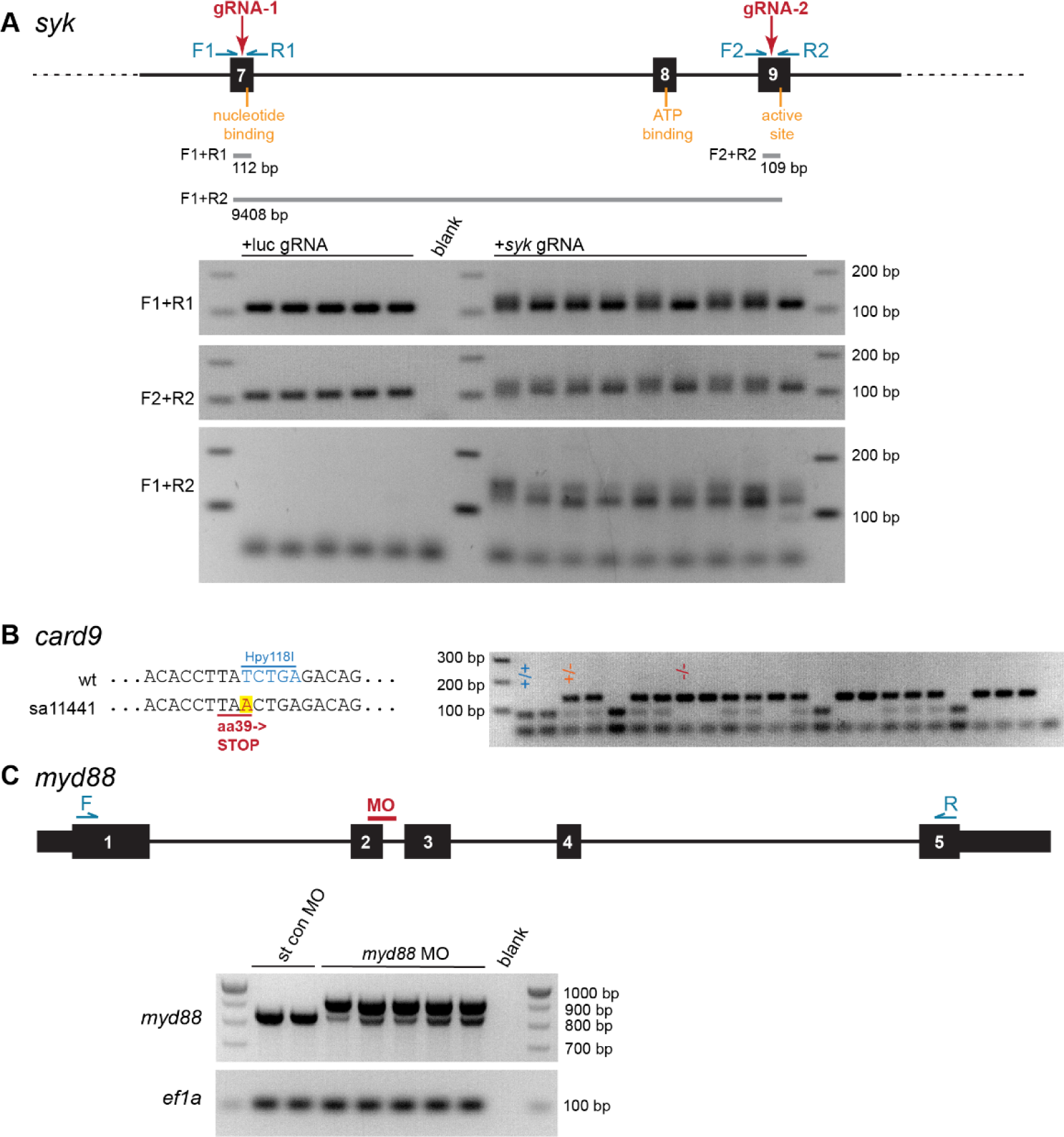
Genetic targeting of *syk*, *card9*, and *myd88.* **A.** Schematic of gRNA target sites in *syk* gene and example DNA gels showing PCRs performed to confirm successful targeting. **B.** Schematic of single point mutation present in *card9* sa11441 mutant allele and example DNA gel showing PCR and restriction digest performed to genotype zebrafish. **C.** Schematic of morpholino target site in the *myd88* transcript example DNA gel showing RT-PCR performed to confirm successful splice blocking.

**Supplementary Figure 2.**
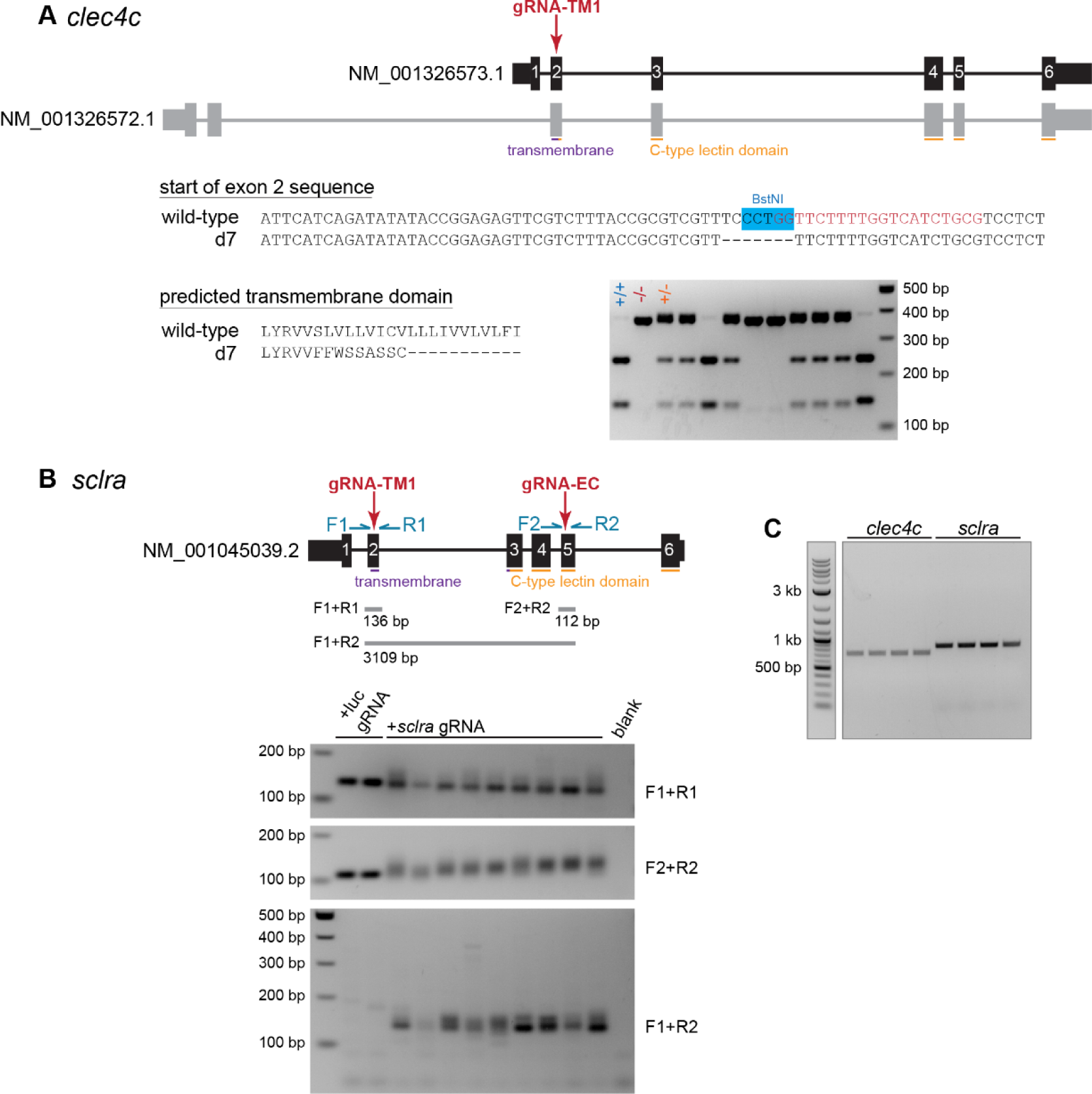
Genetic targeting of *clec4c* and *sclra.* **A.** Schematic of gRNA target site in *clec4c* gene and putative RefSeq transcripts, DNA sequence of start of exon 2 in isolated d7 mutant line, protein sequence of predicted transmembrane domain in isolated mutant line, and example DNA gel showing PCR and restriction digest performed to genotype zebrafish. **B.** Schematic of gRNA target sites in *syk* gene and putative RefSeq transcript and example DNA gels showing PCRs performed to confirm successful targeting. **C.** DNA gel showing amplification of *clec4c* and *sclra* transcripts from zebrafish larvae by RT-PCR. Gradient PCR was done, each lane shows amplification with a different primer annealing temperature.

**Supplementary Figure 3.**
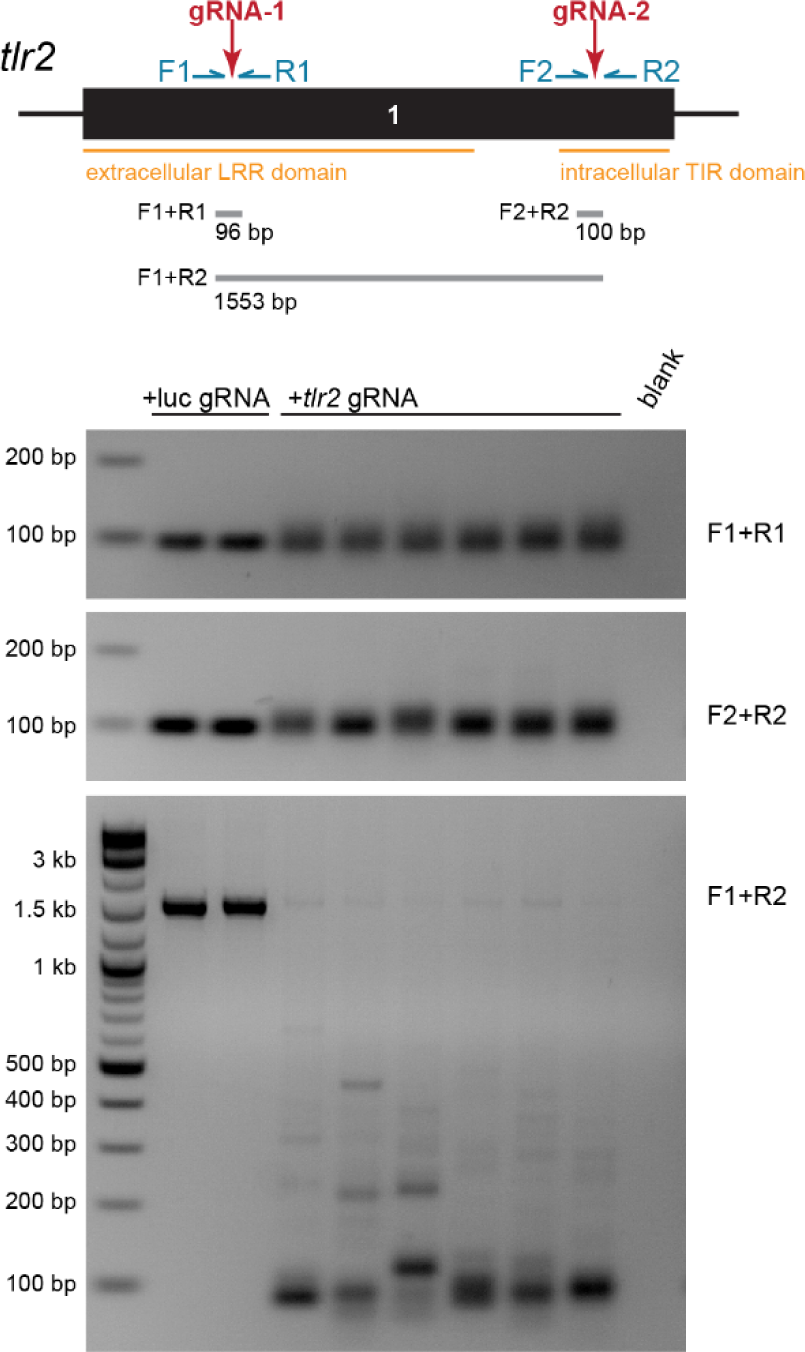
Genetic targeting of *tlr2.* Schematic of gRNA target sites in *tlr2* gene and example DNA gels showing PCRs performed to confirm successful targeting.

